# Drivers of change and ecosystem status in a temperate lake over the last Post-Glacial period from 14.5 kyr

**DOI:** 10.1101/2020.03.27.011502

**Authors:** I Tõnno, R Freiberg, L Talas, A Kisand, S Belle, N Stivrins, T Alliksaar, A Heinsalu, S Veski, V Kisand

## Abstract

Understanding the long-term dynamics of ecological communities on the centuries-to-millennia scale is important for explaining present-day biodiversity patterns. Placing these patterns in a historical context could yield reliable tools for predicting possible future scenarios. Paleoarchives of macro+ and micro-fossil remains, and most importantly biomarkers such as fossil pigments and ancient DNA present in various sedimentary deposits, allow long term changes in ecological communities to be analysed. We use recent compilations of data including fossil pigments, metabarcoding of sedimentary ancient DNA and microfossils together with data analysis to understand the impact of gradual *versus* abrupt climate changes on a lake`s ecosystem status over the last 14.5 kyr. We give examples of hypotheses that need long-term data, which can be addressed using well-established paleoproxy variables. These variables describe the climate, together with vegetation change and the appearance of anthropogenic forcing, either as a gradual change or an abrupt event. We were able to detect abrupt changes in the lake ecosystem during the stable period of the Holocene Thermal Maximum and we highlight the increased frequency and degree of perturbation in lakes due to non-immediate human activity over a larger region. Both observations demonstrate an impaired relationship between a gradual external driver and ecosystem response and apply to future scenarios of climate warming and increased human impact in north-eastern Europe.

## 1. Introduction

Paleoecology in general has proved our ability to use long term data to test ecological theories, hypotheses, and climate models that need data over long timescales with broad taxonomic coverage. Almost all ecosystems are under constant pressure from changing climatic drivers, which can be gradual or abrupt [1,2]. Not only abrupt change in drivers but also gradual changes can ultimately cause abrupt (threshold-like) responses in ecosystems [3,4]. Lakes are usually closely influenced by the surrounding catchment, so the response of their ecosystems to outer physical (i.e. climate change) or anthropogenic forcing could be indirect, mediated through processes in the catchment [5,6]. Current climate change is a rapid process, equally prone to cause immediate abrupt changes and to pave a way for much faster breakdown of the resilience of ecosystems than any event in the recent past [7,8]. Such breakdown can be facilitated by a combination of anthropogenic forcing and climate change, e.g. eutrophication and soil erosion increasing together because of human activities [9]. The term ‘resilience’, common in contemporary ecology, is becoming more and more popular in paleoecology studies (reviewed in [10]) and means the ability of a system to retain its current state via tolerating or resisting disturbance without rearranging into a structurally different state.

Detecting past regime shifts in paleoecological data series enables the response pattern of ecosystems to gradual and abrupt changes to be deciphered. In addition, reconstruction of past changes in ecological states and variability of ecosystem characteristics during stable climatic periods (or gradual successions with gradual responses) can be elucidated [4,11]. Synthesis of such knowledge provides potential warning signs for future scenarios [12]. Disentangling how lakes responded to climate change (including warming) in the past is important, because according to prevailing scenarios the global temperature will increase by up to 6 C° by the end of the 21^st^ century [7].

Primary producers are the base of most aquatic food webs, therefore variability in their biomass and productivity leads to predominant control over the whole lacustrine ecosystem [13,14]. Several primary producers in a lake are short–living benthic or pelagic algae that can respond rapidly to changing environmental conditions such as water temperature, humification and nutrient load [15–18]. Naturally, the lake responds to a warmer climate with increased water temperature, so thermophilic bloom-forming cyanobacteria could gain a competitive advantage over other algal groups in the near future [19,20] in temperate regions. However, there is no basic information on how total species richness and diversity will change. Climate warming has occurred previously in Northern Europe, e.g. there were several abrupt changes in the Post-Glacial period [21,22] followed by more gradual changes over the most of the Holocene [23,24], allowing a relationship between forcing and ecological responses to be identified with predictive power. However, there are still gaps in understanding of the causal relationships between past climate changes and ontogenic development in lakes. We studied fossil pigments from Lake Lielais Svētiņu (hereafter LS) in Eastern Latvia as a proxy for the whole community of primary producers. The lake itself was in the natural state throughout the Holocene with relatively late and gradual human intervention [25,26]. However, a direct (~2.0 kyr) human footprint is clear and important enough in the area, making LS a useful model for studying natural and human-related external forcing of aquatic ecosystems. Previous paleoresearch on LS has focused on phytoplankton composition using non-pollen palynomorphs [26–28], ancient sedimentary DNA (sedaDNA) [28,29], sediment organic matter (OM) and geochemical composition [30], while energy flows in food webs were studied by the stable isotopic composition of subfossil chironomids (Diptera, Chironomidae) [31]. Furthermore, Veski et al. [21,32] and Stivrins et al. [25,26] used pollen data from LS sediments to reconstruct past climate and vegetation history in the eastern Baltic area within the Late-Glacial and Holocene periods. The accumulated data about this lake enable stronger generalisations to be made and allow ecological theories about the response of lacustrine ecosystems to climate change to be tested.

The aim of this study was to identify the response of the primary producer community in a temperate lake to changing external forcing over the last ~14.5 kyr. Using statistical modelling, we considered model fitting and statistical anomalies as indicating that ecological thresholds are approached (perturbation periods and change points). In model fitting, the dynamics of fossil pigment concentration was modelled as a response variable to climate (i.e. temperature), pollen and non-pollen microfossil-based paleoecological proxies and richness of taxon-like molecular operational taxonomic units (mOTUs) from sedaDNA. We hypothesised that although underlying gradual climate change causes mostly smooth successions in algal communities, and strong abrupt climate changes are reflected by abrupt responses, threshold-type ecological changes (regime shifts) have occurred several times over the Post-Glacial period in lakes of Northern Europe. These change points were responses to gradual forcing. In addition, we hypothesised that the richness and diversity of algal organisms change most rapidly during complex forcing periods, e.g. due to anthropogenic impact.

## 2. Materials and methods

### (a) Study site

LS (mean depth 2.9 m; maximum depth 4.9 m) is a drainage lake in the Eastern Latvian lowland with a current area of 18.8 ha and an altitude of 96.2 m above sea level (56°46’N; 27°08’E; Fig. 1). Nowadays the lake belongs to the mesotrophic-dystrophic type (brownish humic water) and has three small inlets and one outlet. Its catchment area (~19.7 km^2^) is mostly forested with some fields south-east of the lake and peatlands in the north-western part with relatively modest and recent human impact [25,30]. Although Latvia has been inhabited by humans since the Paleolithic, the continuous presence of forests in the vicinity of LS verify that this boggy area was not a suitable location for human settlement or activities before calibrated 1500 years [25].

**Figure 1.**
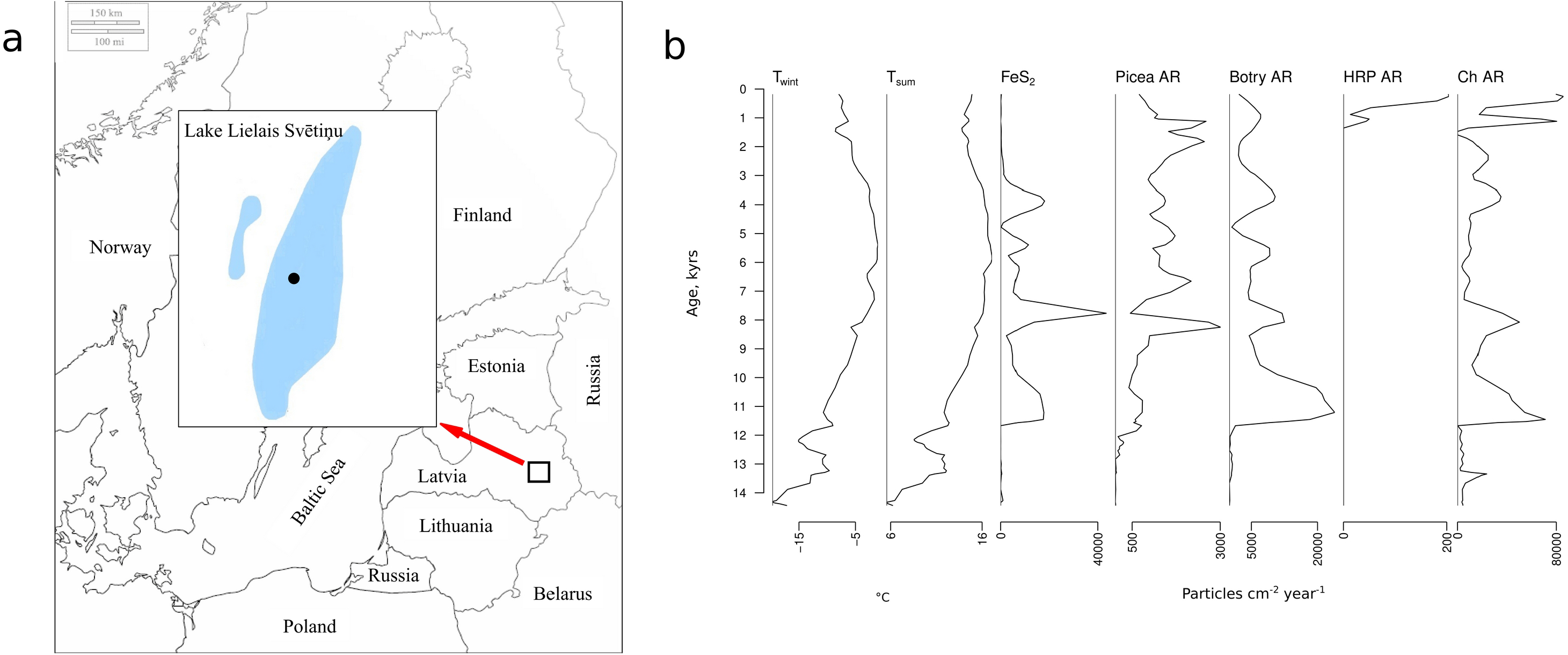
Sampling location, and stratigraphy diagram of basic paleoproxy variables used in the study. Winter temperature – T_wint_; summer temperature – T_sum_; Pyrite – FeS_2_; *Botryococcus* accumulation rate – Botry AR; human-related pollen accumulation rate – HRP AR; charcoal particle accumulation rate – Ch AR

### (b) Coring and chronology

LS was sampled from the ice in March 2009 and March 2013 using a Russian-type peat corer with a diameter of 10 cm. The detailed description and lithology of the 11.3 m long sediment core sampled in 2009 is given in Stivrins et al. [26]. In order to work with fresh sediment material, LS was sampled again for DNA extraction in 2013 and a 10.4 m sediment core was retrieved close to the earlier coring location. Chronology was obtained by cross-correlating lithological changes and loss-on-ignition with the well-dated core from 2009 [29]. The ages in the text are expressed in thousand calibrated years before the present (hereafter, kyr); AD 1950 corresponds to 0 kyr.

Sediment cores from 2013 were sliced into continuous 1 cm thick subsamples and stored at −84 °C prior to analysis. Standard methods were used to determine the OM, mineral matter (MM) and carbonate matter (CM) from sediment subsamples [33]. In the present study, pollen-based reconstructed mean summer temperatures (T_sum_) and mean winter temperatures (T_wint_) from Stivrins et al. [26] were used to characterise the climate as a main driver of the length and intensity of the productive season and as a proxy for possible ice conditions.

### (c) Microfossils and paleoproxy variables

To reveal the combination of environmental changes in the lake ecosystem and the surrounding landscape (regional changes) we used microfossil data and proxy variables calculated using the abundance and diversity of those microfossils. To determine the level of humification of the LS catchment area soils and the lake water, the accumulation rate (AR) of *Botryococcus* (*Botry*) microfossils [26] and the AR of *Picea* pollen [25] were used. The sum of *Secale cereale*, *Hordeum vulgare, Triticum aestivum* and *Avena sativa* pollen ARs (human-related pollens; HRP) from Stivrins et al. [25] was used as a proxy for increased anthropogenic impact. In the present study we used calculated estimates of shade tolerance (S_tol_) and relative openness (R_open_) from Stivrins et al. [26] to characterise the density of the vegetation in the lake catchment area.

Charcoal (Ch) particle AR identified from the pollen slides in Stivrins et al. [26] were used to infer the fire dynamics in region around the lake, which could have influenced the vegetation in the region directly and therefore affected the lake ecosystem indirectly. Pyrite (FeS_2_) from Stivrins et al. [26] was used to indicate anoxic conditions at the bottom of the lake.

### (d) Paleopigment analysis

The resolution of paleopigment subsamples from the 2013 core was one sample (~5 g of wet sediment) every 5 cm. Analysis of the 93 paleopigment subsamples collected followed the recommendations of Leavitt and Hodgson (2001) and is described in detail in the supplemental material. The pigments were separated by reversed–phase high–performance liquid chromatography (HPLC), using a Shimadzu Prominence (Japan) series binary gradient system with a photodiode array (PDA) and fluorescence detector (see Tamm et al. [34] for details). The concentrations of the sedimentary pigments studied were expressed as nanomoles per gram organic matter (nmol g^−1^OM).

Altogether, 12 paleopigments from diatoms, cyanobacteria, chlorophyta, cryptophyta and dinophyta, and those representing total algal abundance and primary production, were separated and identified from the sediment core studied. The affiliation of these paleopigments is reported in detail in the supplemental material. In the present study, the Chl *a*/ Phe *a* ratio was used to follow the preservation conditions of paleopigments in the lake sediments [35]. The ratio of the paleopigments (Chl *a*/ Phe *a* – degradation index – DI) was used in statistical analyses in order to compensate the effect of degradation on the marker pigment dynamics.

### (e) sedaDNA extraction, amplification and sequencing

sedaDNA was extracted from subsamples collected in three biological replicates from each core layer, a total of 252 samples covering the range from 0.05 to 12.0 kyr. All subsamples were collected with 1 ml sterile syringes from the middle of the core. To avoid possible contamination, only the second half entering the syringe was placed in a sterile 2 ml Eppendorf tube. All samples were taken under a positive-flow hood and the subsamples collected were stored at − 84 ̊C in an ancient DNA dedicated lab.

Phytoplankton richness was determined using PCR-based amplicon sequencing (Illumina) of the universal 18S rDNA gene fragment [29], and the ITS2 region [36] was used to assess fungal richness in the lake sediments. Quality-checked raw reads were paired and clustered together (97% similarity threshold) using VSEARCH to receive the mOTUs.

Ecological roles were assigned to fungal mOTUs on the basis of ITS2 sequences to determine the possible relationship between the algae and fungi identified. Fungal trophic state, lifestyle and host groups were determined using information from FUNGild [37] and from relevant publications (e.g. [38–42]. All fungal mOTUs described as possible or obligate parasites of algae, hyperparasites of zootrophic fungi on phytoplankton, parasites of zoo- and phyto-plankton, or found connected to algal hosts, were assigned to the group “possible algae parasites”.

### (f) Statistical analysis

Principal component analysis (PCA) was used for unconstrained ordination of fossil pigment concentrations, and redundancy analysis (RDA) was used to relate paleoproxy variables to the pigment concentration ordination space. RDA uses permutations to test the strength (pseudo-R^2^) and significance (permutation p-value) of each of the explanatory variables in relation to the ordination space.

General additive modelling (GAM) was used as a continuous-time, first-order autoregressive process to account for temporal autocorrelation to describe the statistical relationship between paleoproxy variables, or richness of algal and fungal mOTUs, and PC1 and PC2 scores. PCA scores were considered as proxies for the temporal dynamics of fossil pigment concentrations. The fitted values of GAMs were also compared visually with observed PC score values to reveal non-predictable periods due to gradual changes in climate and catchment conditions. We expected to find a greater difference between model predictions and the observed PC scores during periods of perturbation, which are characterised by more frequent regime shifts or change points; they were obtained from Bayesian model selection, which identifies the strongest association of the linear regression splines of covariates [43]. Thereafter, Bayesian change point analysis was applied to the PC1 scores, as the primary component depending on climate change, to detect abrupt changes in the temporal variability of fossil pigment concentrations.

The statistical software R [44] was used for all statistical analyses with the packages “vegan” [45] for ordination (PCA and RDA), “mgvc” [46] for general additive model (GAM) calculations and “bcp” [43] for Bayesian change point analysis.

## 2. Results

### (a) Stratigraphy of fossil pigments

DI fluctuated on a large scale during the Late Glacial indicating variable preservation conditions during this period (Fig. 2). The contents of the paleopigments investigated and their ratio to DI (hereafter DI ratio) during the Late Glacial (~14.5 – 11.65 kyr) were relatively low and fluctuated somewhat, with a considerable increase during the Allerød warming period (~13.3 – 12.7 kyr) and a sharp decrease thereafter during the Younger Dryas cooling (~12.7 – 11.65 kyr; except Cantha). Within the Holocene period, DI steadily decreased from the older to the younger sediment layers. During the Early Holocene, ~11.65 to ~7.5 kyr, the content of some of the markerpigments studied (D+D, Zea, Lute, Allo) and their DI ratios gradually increased while the contents of Cantha, Echi, Chl b, Perid and their DI ratios remained relatively stable (Fig. 2). The values of the studied paleopigments and their DI ratios were maximal ~6.0 kyr, in the Mid Holocene. Thereafter, the contents of paleopigments and their DI ratios decreased, followed by another increase after ~2.0 kyr, except in Chl a and β–car.

**Figure 2.**
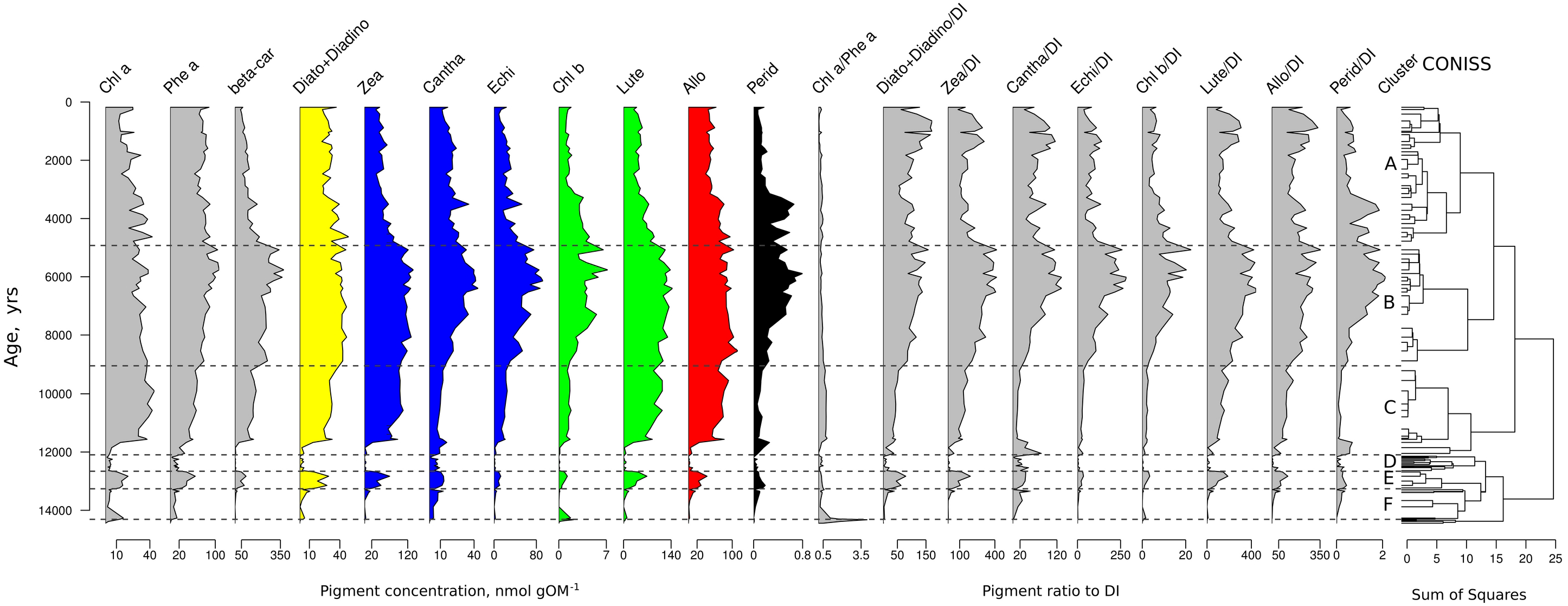
Oscillation of fossil pigment concentration and ratio to degradation index (DI). Chlorophyll *a*–Chl *a*; Pheophytin *a*– Phe *a*; β, β–carotene–β–car; Diadinoxanthin + Diatoxanthin–D+D; Zeaxanthin–Zea; Canthaxanthin–Cantha; Echinenone–Echin; Chlorophyll *b*–Chl *b*; Lutein– Lut; Alloxanthin–Allo; Peridinin–Peri

### (b) Richness of eukaryotic algae in sedaDNA

In total, 257 eukaryotic algal mOTUs were identified over the whole sediment core, the richest phylum being Chlorophyta followed by Chrysophyta, Dinoflagellata and Diatoms.

The richness of algae showed no clear trends until the last 2.0 to 2.5 kyr, ranging from a few to several tens of individual mOTUs per analysed layer. The overall maximum number of mOTUs per layer was 69, with mean = 10. During the last 2.0 kyr there was a trend of increasing richness. The main difference between the last ~2.0 kyr and the earlier Holocene periods was the greater richness of Chlorophyta in the older samples. The richness of all other phyla was greater during the last ~2.0 kyr.

### (c) Richness of possible fungal parasites of eukaryotic algae

The ITS2 analysis enabled a total of 180 mOTUs of fungi possibly parasitic on algae to be detected in the whole sediment core. Most of the possible fungal parasites of algae identified belonged to the phyla *Rozellomycota* (162 mOTUs) and *Chytridiomycota* (17 mOTUs). While most mOTUs were affiliated with a low taxonomic level, some mOTUs could only be assigned at high levels. The greatest richness of fungi possibly parasitic on algae was discovered in the last 2.0 kyr with a maximum of 50 mOTUs per layer (mean 20 mOTUs). The mean richness in the Late Holocene (~4.0–2.0 kyr) was six mOTUs per layer with maximum of 12. The richness in all other periods (Mid Holocene and Early Holocene) was very low (mean two mOTUs and maximum five per layer). Most of the mOTUs of fungi possibly parasitic on algae detected (88%) were located in the last 3.5 kyrs (*Rozellomycota* and *Chytridiomycota*). However, the only species detected in the phylum *Ascomycota* positioned in the Mid Holocene (~4.7 kyr).

### (d) Environmental drivers, disturbance periods and change points

Most of the variation in PC space of fossil pigment concentration is associated with PC1 (85%). The RDA related this variation to major climate and climate-dependent vegetation changes in the region (Table 1, Fig. S1). This reverse was distorted around 2.5 kyr, associated with HRP AR and increased Ch AR in the RDA.

**Table 1.** Summary statistics of redundancy analysis (RDA) using fossil pigment data as dependent variables and various proxy variables as explanatory data. Significant explanatory variables (organic matter – OM; relative openness – R_open_; pyrite – FeS_2_; summer temperature – T_sum_; human related pollen accumulation rate – HRP AR; charcoal particle accumulation rate – Ch AR) were selected using a stepwise procedure (ordistep).

| Final Model | AIC | F | p |
| --- | --- | --- | --- |
| OM | 87.8 | 1.6 | 0.070 |
| $R_{open}$ | 89.1 | 3.0 | 0.055 |
| $FeS_2$ | 89.3 | 3.2 | 0.055 |
| $T_{sum}$ | 90.1 | 4.0 | 0.040 |
| HRP AR | 91.0 | 4.8 | 0.035 |
| *Picea* pollen AR | 93.7 | 7.5 | 0.010 |
| Ch AR |  |  | 0.005 |
|  | 98.1 | 12.0 |  |

To address the potentially non-linear response of algal biomass (and the lake ecosystem in general) to the climate proxy of T_sum_ and other paleoproxy changes, a general additive models (GAM) approach was applied to the PCA sample scores of individual pigments/DI. PC1 is strongly related to T_sum_ (Table 2, model A), while PC2 is related to Ch AR, FeS_2_, and humification proxies (i.e. AR of *Botry,* GAM 2). Comparing the fitted predictions of PC1 to real scores, three disturbance periods deviating from gradual change were identified when GAM A explained the observed dynamics of algal biomass poorly (Fig. 4); prediction of PC2 scores was better using GAM 2 (Table 2, model B). To confirm the association of perturbation periods with possible regime shifts, PC1 score dynamics was subjected to Bayesian change point analysis (Fig. 4b).

**Table 2.** Summary of statistics for the general additive models (GAM) fitted with a Gaussian distribution to PCA scores using fossil pigments (D+D, Zea, Cantha, Echi, Chl b, Lute, Allo and Perid), PC1 with model 1 (A) and PC2 with model 2 (B), respectively. Significant response variables were summer temperature (T_sum_), shade tolerance (S_tol_), charcoal particle accumulation rate (Ch AR), *Botryococcus* accumulation rate (Botry AR) and pyrite (FeS_2_). For fossil pigment abbreviations see Fig 2 legend. “*edf*” denotes effective degrees of freedom.

| Final Model | Terms | edf | F | p |
| --- | --- | --- | --- | --- |
| A. |  |  |  |  |
|  |  |  |  | <0.00 |
| $R^2 = 0.82$ | $s(T_{sum})$ | 6.6 | 7.96 | 1 |
| | $s(S_{tol})$ | 3.9 | 3.01 | 0.0206 |
| B. |  |  |  |  |
|  |  |  |  | <0.00 |
| $R^2 = 0.64$ | $s(Ch\ AR)$ | 6.8 | 3.44 | 1 |
| | $s(Botry)$ | 6.0 | 5.68 | <0.00 |
|  | AR) |  |  | 1 |
|  |  |  |  | <0.00 |
|  | s(FeS<sub>2</sub>) | 5.0 | 8.42 | 1 |

Further, the richness of algae estimated by the number of mOTUs affiliated with pigments/DI of eukaryotic algae was used in the GAMs to explain variation in PC2. Model A (Table 3) indicates that PC2 is significantly related to increased total richness, and specifically to the mOTUs affiliated with Chlorophyta (Table 3, model B). This association is most pronounced for the last 2.5 kyr (Figs. 3 and 4). The non-linear change of richness of parasitic fungi during past millenia was associated most strongly (R^2^=0.479, *p*<0.001) with changes in the potentially additive effect of climate change and human activity.

**Table 3.**
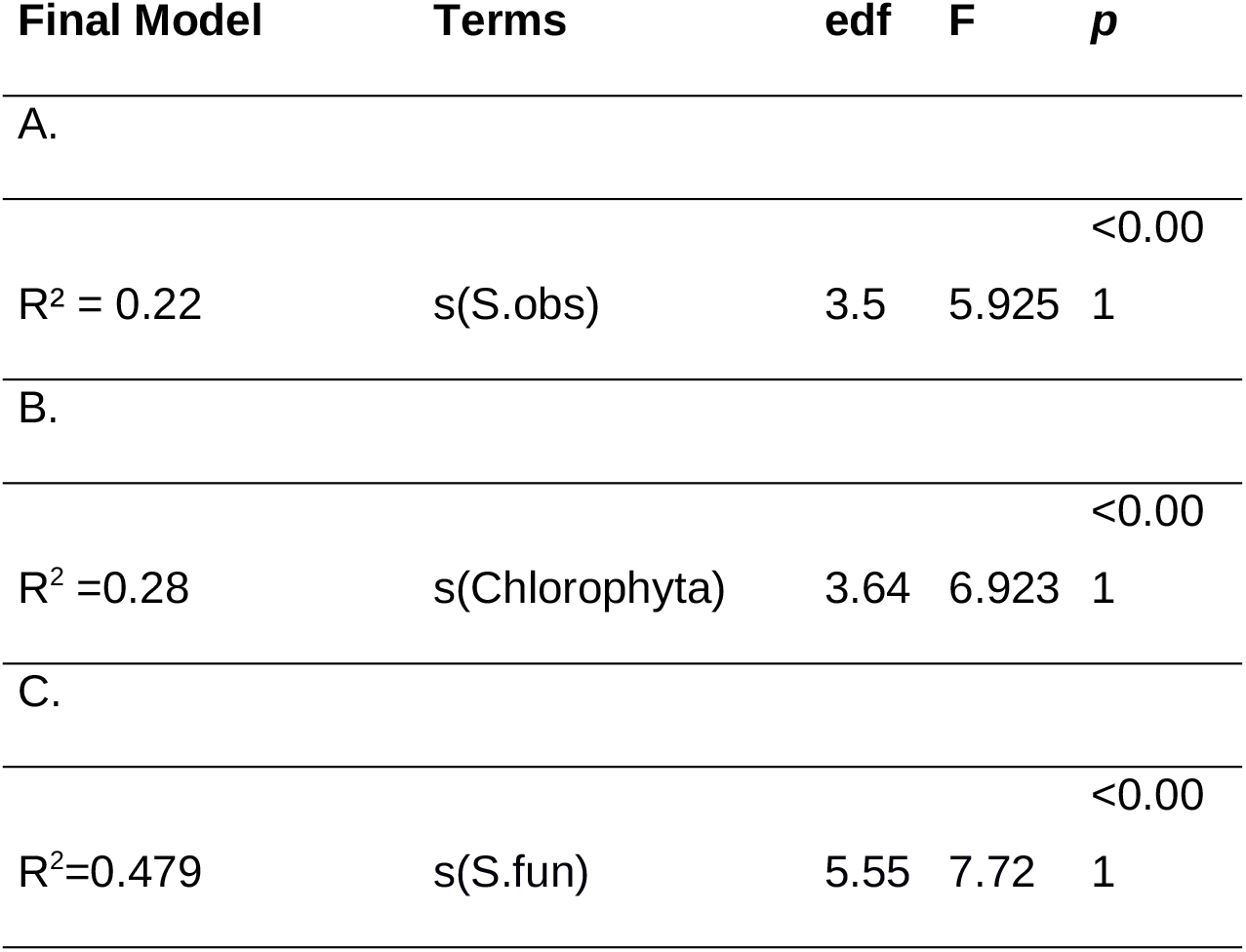
Summary of statistics for the general additive models (GAM) fitted with a Gaussian distribution to PC2 scores using a subset of fossil pigments (excluding cyanobacterial pigments). Total richness of algae (S.obs) (A), richness binned into phyla (B), and richness of fungal parasites/pathogens of phytoplankton (S.fun) (C). In the latter, significant response variables were summer richness in Diatomea and Chlorophyta. “*edf*” denotes effective degrees of freedom.

| Final Model | Terms | edf | F | *p* |
| --- | --- | --- | --- | --- |
| A. |  |  |  |  |
|  |  |  |  | <0.00 |
| R<sup>2</sup> = 0.22 | s(S.obs) | 3.5 | 5.925 | 1 |
| B. |  |  |  |  |
|  |  |  |  | <0.00 |
| R<sup>2</sup> =0.28 | s(Chlorophyta) | 3.64 | 6.923 | 1 |
| C. |  |  |  |  |
|  |  |  |  | <0.00 |
| R<sup>2</sup>=0.479 | s(S.fun) | 5.55 | 7.72 | 1 |

**Figure 3.**
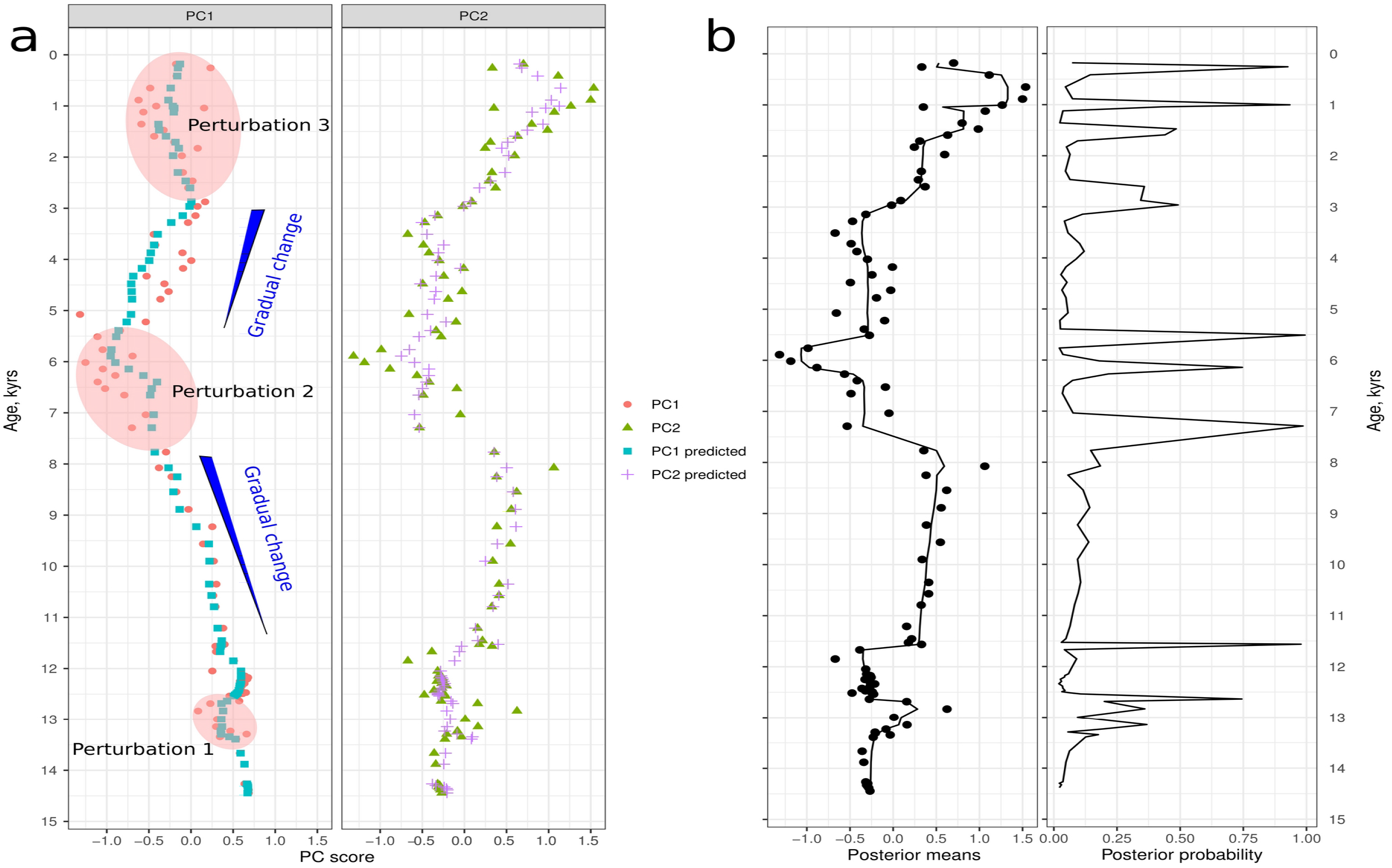
Comparison of dynamics of PC scores of pigment:DI data. (a) Fitted values of these PC scores by GAM using explanatory variables indicated in Table 2. PC1 scores are modelled with T_sum_; PC2 scores with Ch AR and Botry (*Botryococcus*) AR. Blue lines indicate the gradual changes due to temperature change. Red ellipses denote perturbation periods from 1 to 3, which are the time periods used for fitting PC1 to climate (T_sum_) divergence (statistical anomalies) and are better explained by PC2 scores fitted to Ch AR and Botry AR. (b) Bayesian change point analysis of PC2 scores, Posterior Means: location in the sequence versus the posterior means over the iterations (left), location in the sequence versus the relative frequency of iterations that led to a change point - posterior probability (of a change) (right). Indication of regime shifts is highly probable during disturbance periods.

## 3. Discussion

As expected, development of lake ecosystems in the eastern part of northeasten Europe was predominantly modulated by gradual climate changes (i.e. average temperatures) over the period after ice-recession from the area [21,47,48]. The algal community, a keystone functional group of lake ecosystems [14], became gradually more abundant in biomass when both average winter and summer temperatures rose, and most probably formed blooms during the Holocene Thermal Maximum (HTM, ~8.0–4.0 kyr; [18,49]). Thereafter, following climate cooling, the algal biomass decreased. In addition, abrupt temperature changes – strong perturbations to all lifeforms, especially in the Post-Glacial [27] - caused immediate high-amplitude variability in algal biomass. Nevertheless, we were able to identify several perturbation periods leading to change points when the apparently smooth relationship between driver and ecosystem status yielded to a threshold-type response of the algal community and biomass. There were no drastic changes of mOTU richness of algae before ~2.5–3.0 kyr when the increase in richness kicked off. Aquatic fungal mOTUs appeared after the Post-Glacial at low abundance and showed a concerted boost in richness within the Late Holocene. Most of these fungi are parasites of aquatic organisms, primarily algae and zooplankton. This change in organismal richness can be identified as the appearance of a novel external forcing, i.e. anthropogenic pressure. Human impact caused diversification of the eco-niches, leading to higher species richness but also to greater competition (Fig. 4B).

**Fgure 4.**
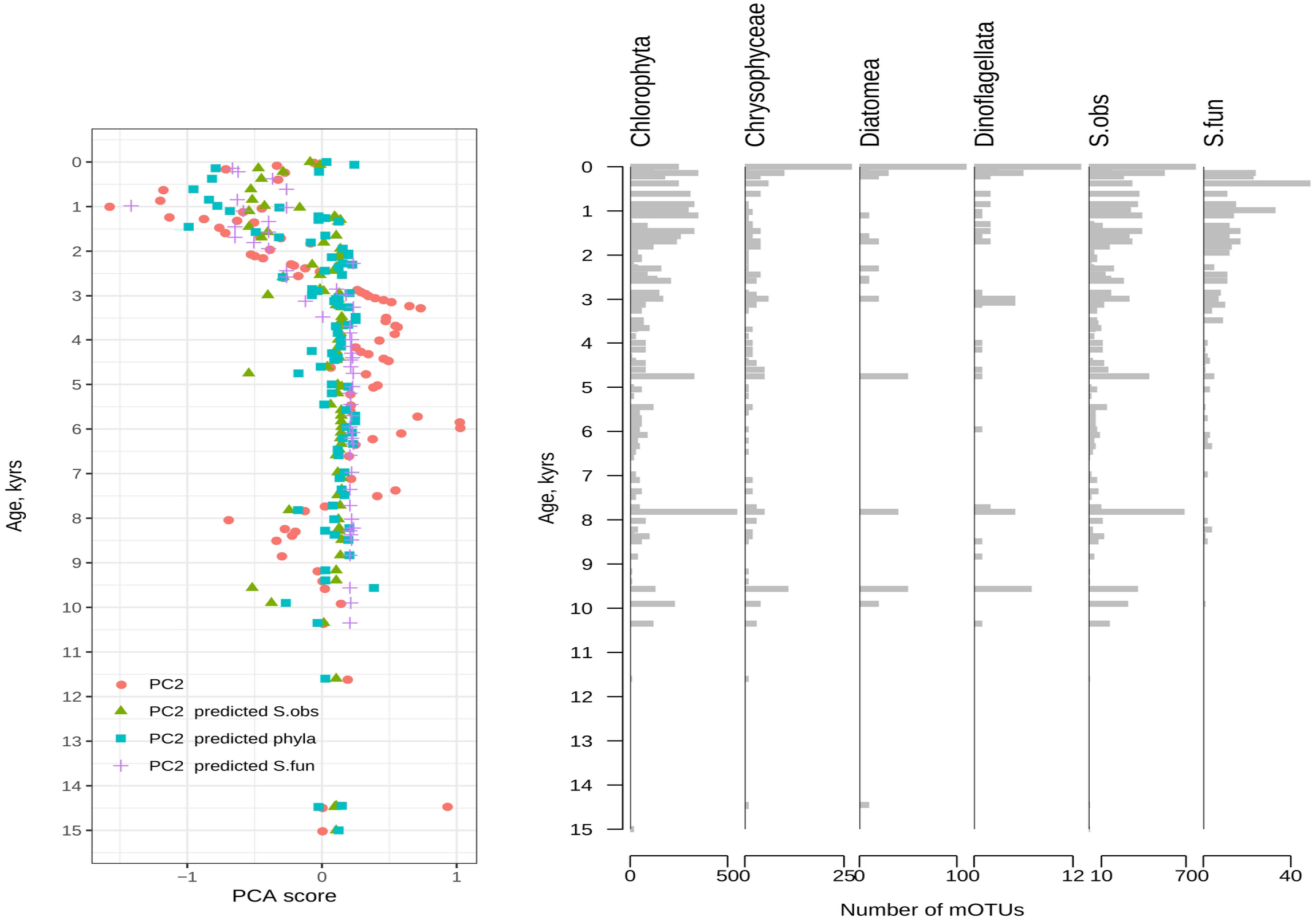
A. Comparison of GAM fitted values with PC2 scores from analysis using the subset of pigment ratios (pigment:DI). Variables used in the modelling are explained in Table 3. B. Stratigraphy of richness of major algal phyla, total richness of algae (S.obs) and fungal parasites/pathogens of algae (S.fun).

### (a) Gradual change periods due to gradual climate change

We observed two major extensive multi-millennia gradual change periods mediated by temperature regime change. Both these periods were in the Holocene. Gradual changes are characterised by readily-predictable dynamics of the PC1 using GAM fitting and Bayesian change point analysis (Fig. 3). From the onset of the Holocene the climate started to warm [21,50], which obviously favoured the increase of algal biomass. This warming also favoured forest expansion in areas surrounding LS [32], clearly reflected in the increase of shade tolerance and decrease in the relative openness indices ([26]; Fig. S2). Changes in sediment composition (increase of OM) and prevailing anoxic conditions (increase of FeS_2_) indicate enhanced production of autochthonous OM, possibly originating from the algal biomass and nutrient-rich allochthonous material inflow to the lake. Indeed, rapid warming favoured terrestrial material inflow to water bodies enriched with nutrients [30]. Surprisingly, no abrupt response could be identified in the present study to the cooling event at ~8.2 kyr [51,52]. The concentrations of some fossil pigments slightly decreased (Fig. 2), but no drastic changes in algal community structure (i.e. perturbation) were observed (Fig. 3b). This can be explained by the possibly high resilience of the algal community, which had enough time to develop into a well-balanced and diverse community as all major mOTU phyla of algae appeared at the beginning of the Holocene (Fig. 4b). The increased algal richness of the lake during the Early Holocene is confirmed by microfossil analysis [26].

The second period of smooth driver–status relationship with a cooling climate (decreasing average summer temperatures and increasing continentality) was shorter, strictly lasting only about 2.5 kyr, from ~ 5.5 to 3.0 kyr. However, low-probability change points were detected up to 1.0 kyr. During this gradual change period, conifer forest developed in areas surrounding LS (revealed by *Picea* pollen and stomata data) and peat decomposition decreased after ~5.0 kyr. The latter observation suggests increased soil moisture levels [26,53], and in combination with shallowing of the lake water due to sedimentary filling processes [31], possibly led to lower light intensity in the lake water column. According to Fritz and Anderson [54], peatland and conifer forest lead to acidification of drainage area soils, which in turn can increase in-lake dissolved organic carbon and decrease the pH. Moreover, the development of conifer forests is generally accompanied by lower soil N, leading to a gradual loss of N from the lake and hampering of in-lake productivity [54]. Such natural climate-driven processes dominated until a new external forcing appeared (see below). The results of the RDA analysis (Fig. S1, Table 1), variability in PC2 (Figs 3 and 4; Table 2) and absorbed covariation of several other environmental proxies (Table 2) revealed an intense association of primary producer dynamics with both in-lake and catchment processes.

### (b) Perturbation periods and their association with change points

Perturbation periods were primarily indicated by the lack of fit between the PC1 scores and the scores predicted by GAM.

During the Late-Glacial, the “young” and less diverse algal community responded abruptly (high biomass versus very low biomass) with no sign of high resilience on the scale of data resolution in the study (interval of a few hundreds of years). Unfortunately, we have no solid mOTU data from the Late-Glacial period. In the few layers we analysed we detected Chlorophyta, Chrysophyta and Diatomea using 18S rDNA fragment analysis. Owing to the scarcity of DNA in these samples, no fungal ITS2 analysis was performed. However, an earlier study using 18S rDNA analysis indicated that fungi were present during the Post-Glacial period [29].

More striking is a perturbation during the HTM, which is considered a stable period in northeastern Europe in respect of the surrounding terrestrial ecosystems [55]. In the algal community there was a relatively high variability of fossil pigment concentrations from around 7.5 kyr to 5.0 kyr (Fig. 2). The change points most probably fall into the same period (Fig. 3), though there is a stable period between the first and second change points lasting over 0.5 kyr. Proposed triggers of the environmental processes behind this perturbation are humification and contrasting ice-conditions – as a single driver for different change points, or a combination of these processes. Humification in LS was already beginning from the early Holocene (Fig. S1). Indeed, the acid water-tolerant genera cryptophyta (Allo) and dinophyta (Perid) became relatively more abundant than most planktonic cyanobacteria (Fig. 2; [56]). This is supported by the appearance of several low pH-tolerant green algae species among the mOTUs affiliated with the genera *Scherffelia, Choricystis, Desmodesmus* and *Woloszynskia* (Fig. S5) [57]. Ice-conditions are directly related to climate conditions (average winter temperatures); the HTM is a period when weak or missing ice-conditions could occur (average T_wint_ around −1.5 to −1 °C). Ice cover conditions respond to small changes in temperature in a threshold fashion [58], so relatively subtle temperature changes can cause dramatic changes in the lake state. A shorter ice cover duration (no light shading snow on top of the ice) can cause a prominent increase in the length of the growing season [59,60], while even a tiny recession from the trend could form the basis for the change points during 7.5–5.0 kyr.

It has been shown that rapid threshold-like responses to increased external driver force are often anthropogenic [4,11]. Direct anthropogenic eutrophication has occurred during the past few centuries. The first minor human farming activity in the landscape immediately surrounding LS started ~2.0 kyr (Fig. 1b, appearance of HRP) with intensification ~0.6 kyr [26,61]. Despite the relatively late immediate anthropogenic forcing, a modest effect clearly coinciding with the increase of R_open_ (Fig. S2) can be assumed in the region (several thousand km^2^) after ~3.5 kyr [62], best described by the dynamics of PC2 on fossil pigments. This is mirrored by several observations: a prominent increase in algal species and their fungal pathogens (number of mOTUs as species proxy, Fig. 4B, Table 3), and an increased frequency of change point posterior probabilities (Fig. 3b). There has been a similar increase in the number of species and diversity due to modest human activity, which we call generic human impact, in large scale areas of vegetation [63–65] and overall richness in the Eukaryote domain [29] in lakes of northeastern Europe. Although the modest human activity increased the patchiness of the boreal and semi-boreal forest, offering more niches for various terrestrial species, the similar increase in the lake’s ecosystem is intriguing. Even more fascinating is the concerted rise of fungal parasites of algae. While the biomass and productivity of the algal community has crushed below the level attained during the HTM, the increase in species richness means capturing new niches or greater competition between algal species for the same resources. Because of such competition leads to suboptimal growth conditions, the increased parasite attack rate is obvious ([66] and references therein).

## 4. Conclusions

Gradual climate change and gradual development of ecosystems have worked together to shape the aquatic ecosystems of north-eastern Europe. Gradual climate warming or cooling can induce natural eutrophication or oligotrophication processes in the lake. Threshold-like responses occur when climate, lake ontogeny, or novel forcing such as human influence disrupt in-lake algal community and climate relationships. The latter can be modulated as generic influence in larger (thousands of km^2^) regions, or direct forcing via traditional agricultural activity near the lake. Modest disruption can lead to increased microbial biodiversity including various pathogens, which could serve as a signature of future perturbation [67] leading to future changes in ecological status.

## Supporting information

Full supplemental material

## Data accessibility

Raw sequencing data were deposited at the European Nucleotide archive under project PRJEB17781.

All metadata are available from the Dryad Digital Repository: (will be deposited when accepted for publication).

## Author’ contributions

V.K. and I.T. conceived of the project. I.T. A.H., S.V. and V.K. collected samples. R.F. and I.T. carried out pigment analyses. S.N., A.K., T.A and S.V. carried our conventional paleoecological analyses. L.T. A.K. and V.K. carried out molecular analyses. V.K., I.T. and S.B. analysed data. V.K. and I.T. prepared the first draft, and all authors contributed to drafts.

## Acknowledgements

The authors acknowledge Dr Anne Bjune from University of Bergen and Prof Heikki Seppä for their general comments on the study design.

