## Supplementary material for "Drivers of change and ecosystem status in a temperate lake over the last Post-Glacial period from 14.5 kyr": Full supplemental material

Study site

Lake Lielais Svētiņu bedrock consists of Devonian dolomite covered by Quaternary deposits with a thickness of 5–10 m consisting of peat, sand, silt, clay and till. The deposits have been greatly paludified during the Holocene [1]. According to our present knowledge, there is no monitoring data available about the nowadays water chemistry and phytoplankton composition dynamics of LS. The climate in the area is a combination of continental (Eurasia) and maritime (Atlantic Ocean), therefore the annual frequency of arctic and sub-polar air masses is fairly high [2].

Coring and chronology

From sediment core sampled in 2009 eighteen radiocarbon dates were obtained from terrestrial remains or bulk sediment by accelerator mass spectrometry analyses performed at the Poznan Radiocarbon Laboratory (Poland) and Tallinn University of Technology (Estonia).

Sediment cores from 2009 and 2013 were documented and transported in cool & dark container to the laboratory for further analyses.

From the sediment subsamples the water content was measured by drying the samples to constant weight at 105 ˚C. The sediment organic matter (OM) was applied as the weight loss-on-ignition at 550 ˚C for 4 h while the ignition residue was estimated as the mineral matter (MM). The differences of LOI between 550 ˚C and 950 ˚C for 2 h multiplied by 1.36 was applied for carbonate matter (CM) content measurements [3].

The pollen-based mean summer temperature [4]was reconstructed within Finland-Estonia-Sweden-West Russia-Lithuania pollen-climate calibration set using the weighted averaging-partial least squares regression and calibration procedure [5,6].

Sediment cores covered Late Glacial (~14.5 - 11.65 cal kyr BP) and Holocene (since ~11.65 cal kyr BP) periods. The content of OM remained low in Late Glacial, followed by gradual increase in Holocene up to ~ 5.0 cal kyr BP. Thereafter slight decrease of OM content towards upper sediment layers is detectable (Fig. 1B). The dynamics of MM and CM was generally opposite to OM one, with decreasing trend from bottom to the topmost sediment core (Fig.1B).

Microfossils and environmental parameters

Remains of *Botryococcus* in the sediment point to the presence of incoming dystrophic waters [7], that can be associated with the paludification processes around the LS [4]. Hence, the presence of a peatland and conifers can lead to acidification of catchment soils which can increase dissolved organic carbon fluxes and cause a decrease of pH in a lake [8].

Paleopigment analysis

The frozen sediment samples were first freeze-dried (Heto PowerDry LL3000) 48 h in the dark. Thereafter pigments were extracted with acetone-methanol mixture (80:20 v:v) at -20 ˚C in the dark for 24h. Finally all pigment extracts were clarified by filtration through 0.45 µm filter (Millex LCR, Millipore) to remove any particles. For paleopigment peak identification and quantification commercially available external standards from DHI (Denmark) was used.

**Table S1. Affiliation and abbreviation of studied paleopigments.**

| **Pigment name** | **Abbreviation** | **Affiliation** | **Reference** | **Notes** |
| --- | --- | --- | --- | --- |
| Chlorophyll *a* | Chl *a* | Tot algal abundance and primary production | [9,10] | |
| Pheophytin *a* | Phe *a* | Tot algal abundance and primary production | [9,10] | General derivative of Chl *a* |
| β, β–carotene | β–car | Tot algal abundance and primary production | [9,10] | |
| Diadinoxanthin* | Diadino | Diatoms | [11] | |
| Diatoxanthin* | Diato | Diatoms | [11] | |
| Zeaxanthin | Zea | Tot cyanobacteria | [12,13] | Could be found also in chrysophytes and in green algae |
| Canthaxanthin | Cantha | Colonial & filamentous cyanobacteria | [10] | |
| Echinenone | Echin | $N_2$-fixing filamentous cyanobacteria | [14] | |
| Chlorophyll *b* | Chl *b* | Green algae | [9,10] | Also in higher plants |
| Lutein | Lut | Green algae | [9,10] | Also in higher plants |
| Alloxanthin | Allo | Chryptophyta | [11] | |
| Peridinin | Peri | Dinophyta | [11] | |

* In present study diatoms were represented by complex pigment formed by carotenoids diadinoxanthin and diatoxanthin (D+D) as in their xanthophyll cycle diadinoxanthin could be transformed to diatoxanthin under excessive light [11].

sedaDNA analysis

Total DNA extraction from sediment samples (~0.2–0.3 g wet sample) was done using PowerSoil® DNA Isolation Kits (MoBio Laboratories) under positive-flow hood (Kojair K-safety KR-125). All the surfaces were previously exposed to UV light and were cleaned with Thermo Scientific™ DNA AWAY™ Surface Decontaminant. Extracted DNA samples were stored for short-term at -20℃ or for long-term at 80℃. DNA extraction and following PCR were conducted in separate laboratories.

Richness of phytoplankton was determined using PCR based amplicon of universal 18S rDNA gene fragment - hypervariable region V4 (length of ~300 - 350 bp) according to earlier publication (Kisand et al. 2018). The ITS2 region was used to assess the fungal richness from lake sediments (Tedersoo et al. 2014). ITS2 region amplification was received using multiplex primer pairs (forward primers ITS3-Mix1-tag, ITS3-Mix2-tag, ITS3-Mix3-tag, ITS3-Mix4-tag, ITS3-Mix5-tag and reverse primer ITS4mod-tag) [15]. The amplification of regions (ITS2 region — annealing at 46°C, 18S region — annealing at 52°C) and workstation processes were executed as described by Kisand et al. [16].

The amplicon libraries, usually 48 samples pooled, were purified using PCR Kleen (Bio-Rad), tags added and send to FIMM, University of Helsinki, Finland for Illumina sequencing using the PE250 chemistry.

#### Sequences analysis and mOTU clustering

All demultiplexed raw reads were quality trimmed by removing Illumina-specific sequences and low-quality nucleotides (<Q30) (Trimmomatic V0.36), then paired together and full-length de-replicated. All reads that were <200 bp in length or contained chimeras were discarded from the dataset. Surviving reads were clustered together by (97% of similarity threshold) using VSEARCH to receive mOTUs (molecular operational taxonomic units). For ITS2 dataset the additional post-clustering curation was conducted using R-package LULU [17] to reduce the erroneous mOTUs. To obtain taxonomy, the representative sequences of clusters were compared using BLASTn against UNITE database (UNITE ver 7; [18]). We assigned mOTUs to species, genus, family, order or class level by sequence identity criterion of 98%, 90%, 85%, 80%, 75% [15]. All non-fungal organisms were removed from further analysis.

### Results

Association between phytoplankton biomass, richness and environmental variables

According to paleopigment concentration the phytoplankton biomass followed the climate changes during the whole study period. This is best demonstrated by PCA and RDA (Fig 3).

According to RDA major climate and climate depending vegetation changes were the increase of summer temperature ($T_{sum}$), higher density of vegetation (increased $S_{tol}$ and decreased $R_{open}$), increased proportion of OM, higher humification (accumulation rate of pollen of *Picea* and microfossils of *Botryococcus*) and decreased oxygen (higher amount of pyrite particles in the sediment).

Richness of eukaryotic algae in sedaDNA

Identification of various algae using 18S rDNA fragment does not allow very high resolution of taxonomy, therefore only few (n=3) mOTUs were possible affiliate at genus level, both belonged to *Dinoflagellata* (two *Ceratium* spp and one *Woloszynskia* sp mOTUs), rest of mOTUs remained classified at higher taxonomical resolution.

Richness of possible fungal parasites of eukaryotic algae

According to ITS analysis a very small number of mOTUs were represented by phylum *Ascomycota* (1 mOTU) and *Chytridiomycota* (17 mOTUs).

17 mOTUs were assigned at the order level (*Lobulomycetales* (2 mOTUs), *Rhizophydiales* (9 mOTUs), *Chytridiales* (6 mOTUs) in phylum *Chytridiomycota* and 1 mOTU at the family level (*Halosphaeriaceae*, phylum *Ascomycota*).

**Supplementary material**

Figure S1. Variation of fossil pigment ratio to DI over the last ~14.5 kyr analysed by principal component (PC) analysis: a) PC analysis of fossil pigment:DI (for pigment abbreviations see Fig. 2); b) relationship of fossil pigment:DI with set of significant explanatory variables analysed by RDA (for significant variables and their abbreviations see Table 1). Organic matter–OM (%); relative openness–$R_{open}$; summer temperature–$T_{sum}$; pyrite–$FeS_2$; human related pollen accumulation rate–HRP AR; charcoal particles accumulation rate–Ch AR.

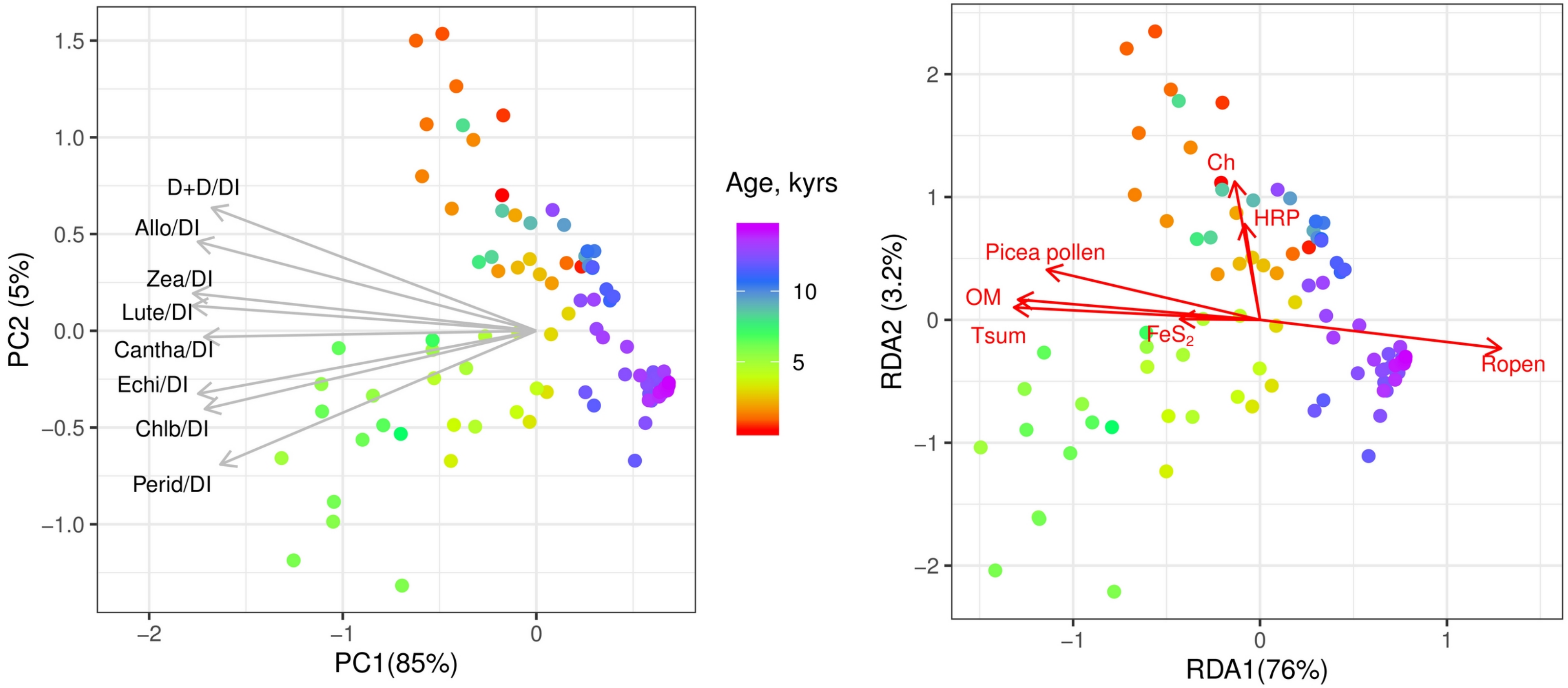


Figure S2. The stratigraphic diagram of PCA scores of fossil pigments, climate (continentality ($T_{sum}/T_{wint}$)–Cont; vegetation proxy variables (shade tolerance–$S_{tol}$; drought tolerance–$D_{tol}$; waterlogging tolerance–$W_{tol}$; relative openness–$R_{open}$), and organic matter–OM (%).

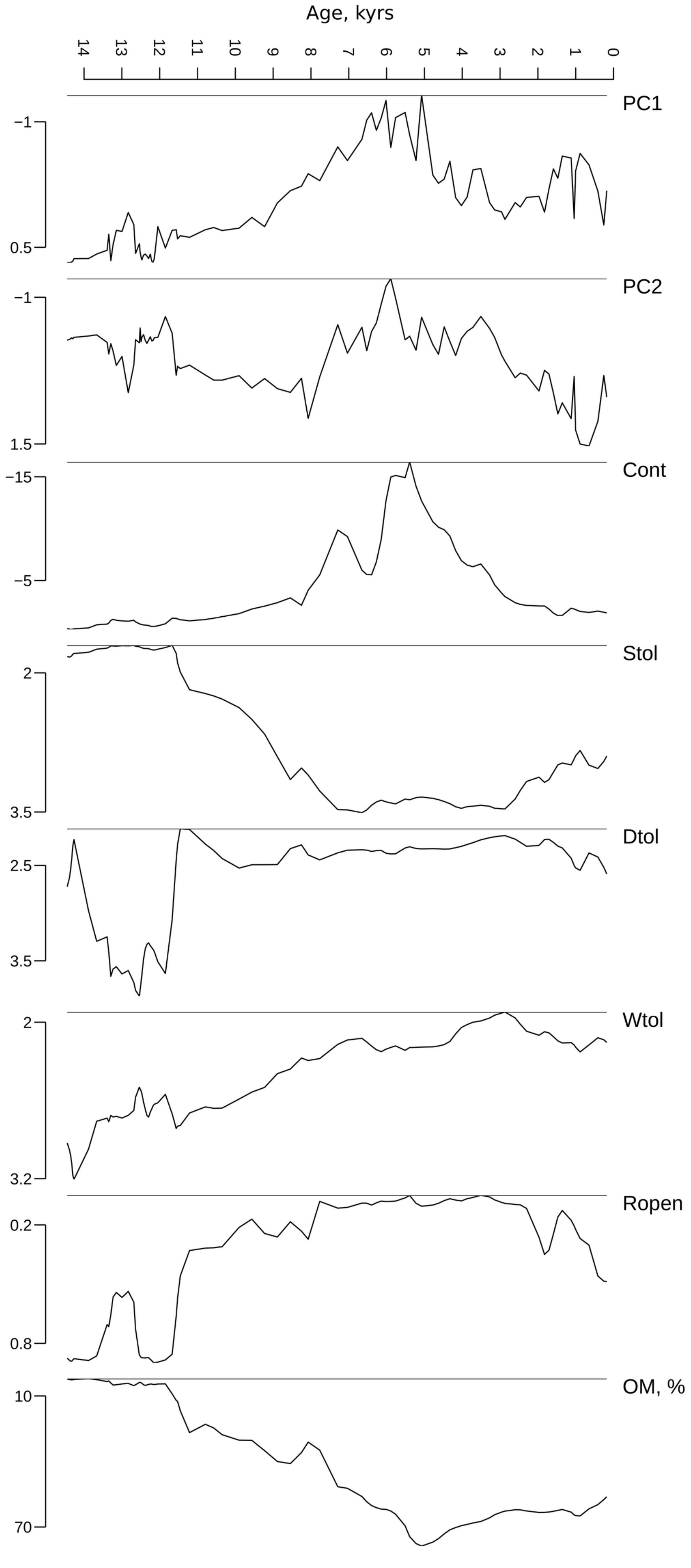

Age, kyrs
14
13
12
11
10
9
8
7
6
5
4
3
2
1
0
PC1
−1
0.5
PC2
−1
1.5
Cont
−15
−5
Stol
2
3.5
Dtol
2.5
3.5
Wtol
2
3.2
Ropen
0.2
0.8
OM, %
10
70

Figure S3a.  Plot of the smooth component of model A (see Table 2)

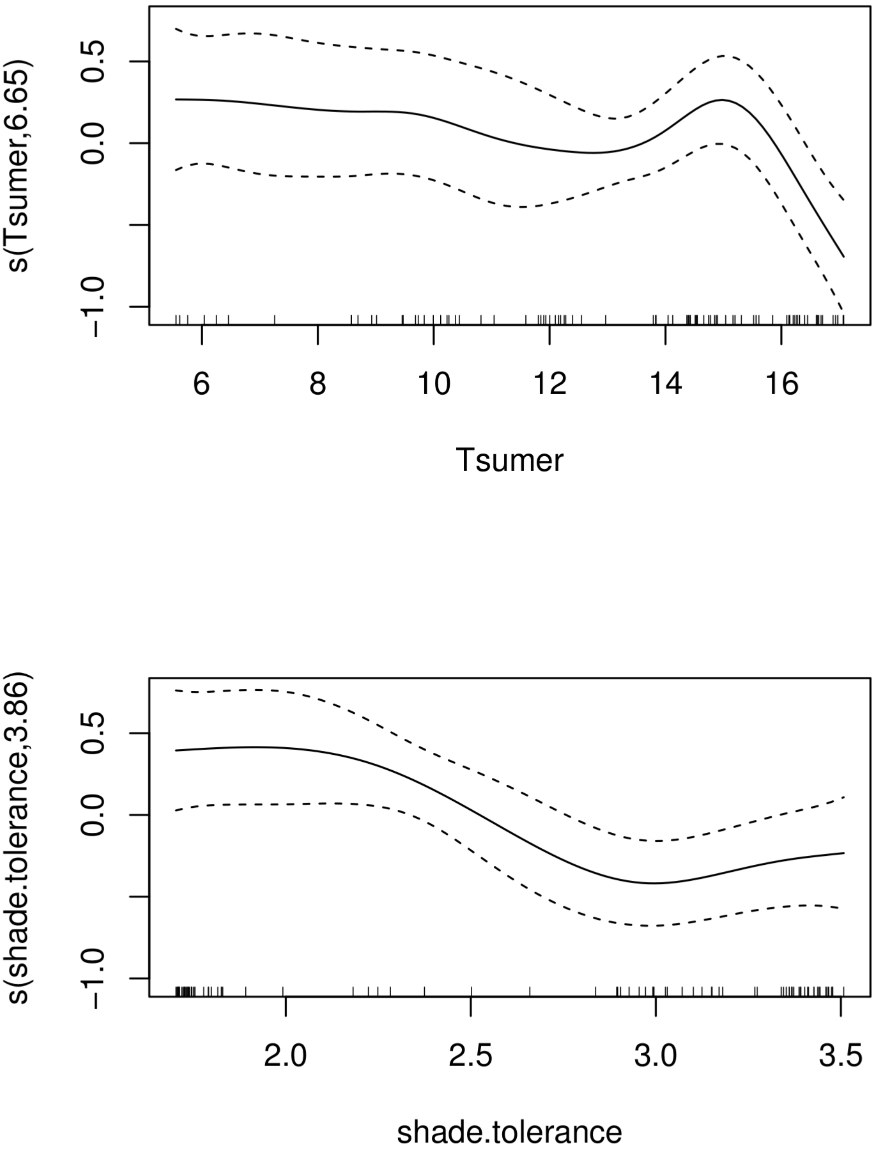

Figure S3b. Plot of the smooth component of model B (see Table 2)

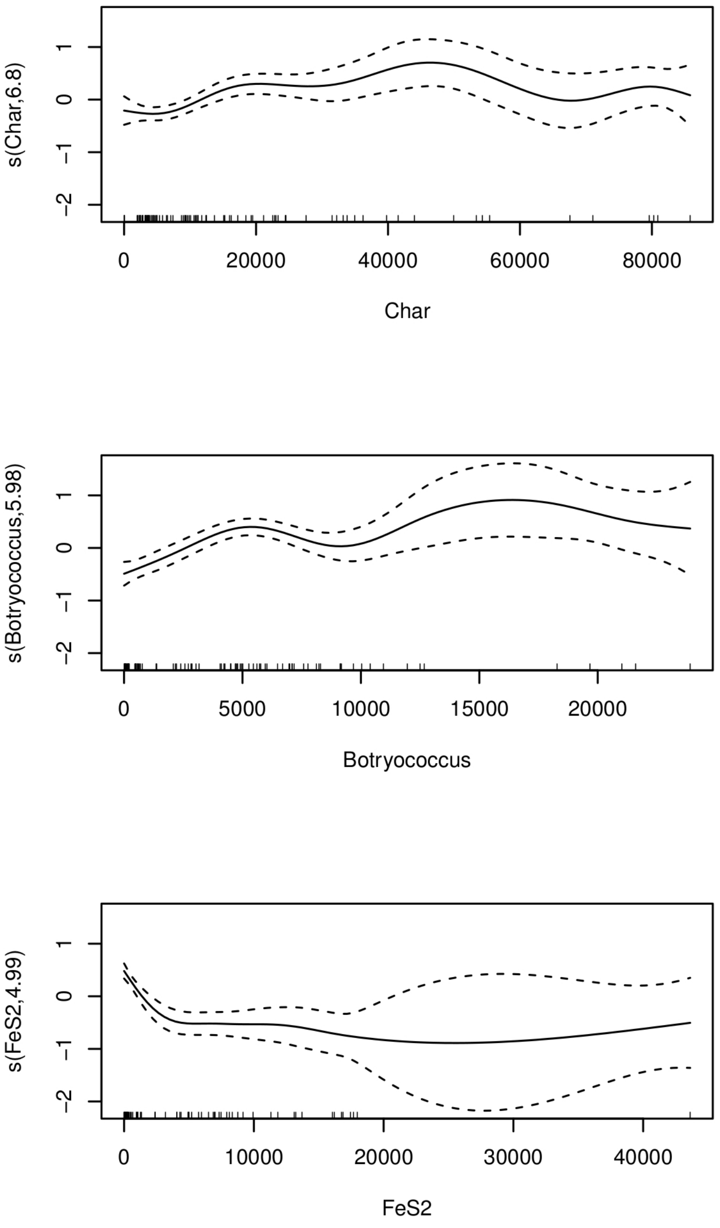

Figure S4a. Plot of the smooth component of model A (see Table 3).

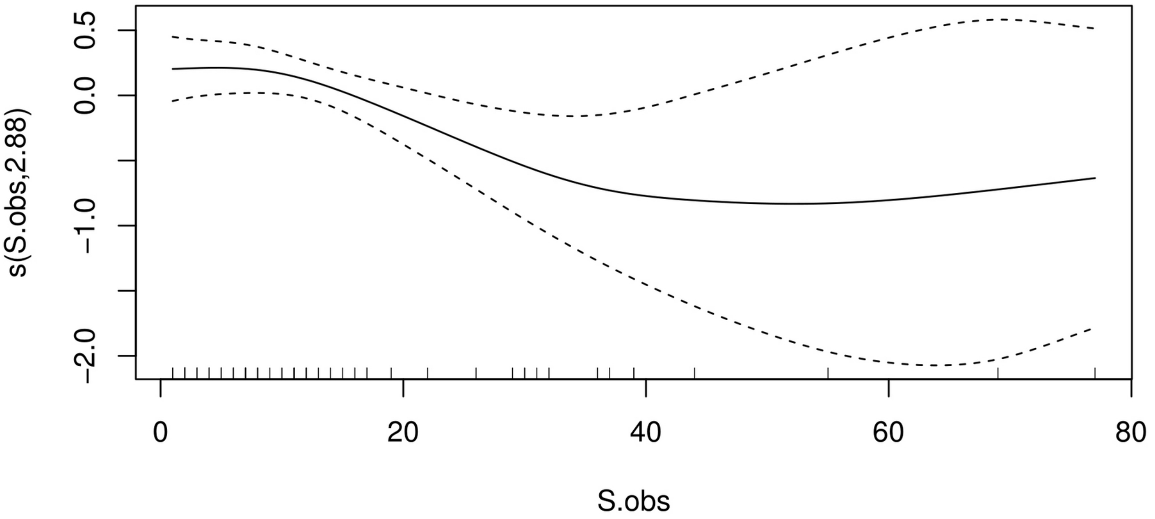

Figure S4b. Plot of the smooth component of model B (see Table 3).

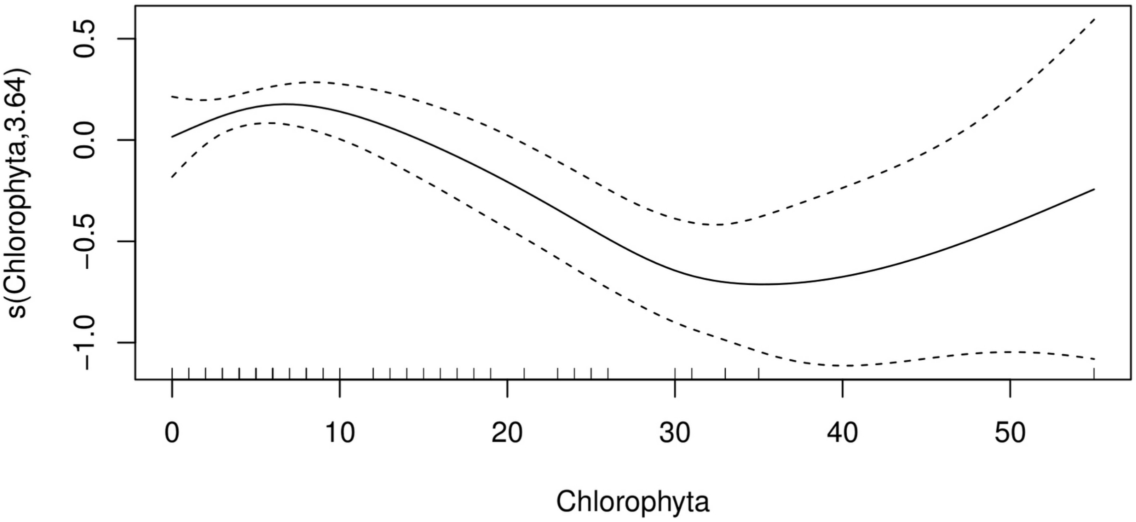

Figure S4c. Plot of the smooth component of model C (see Table 3).

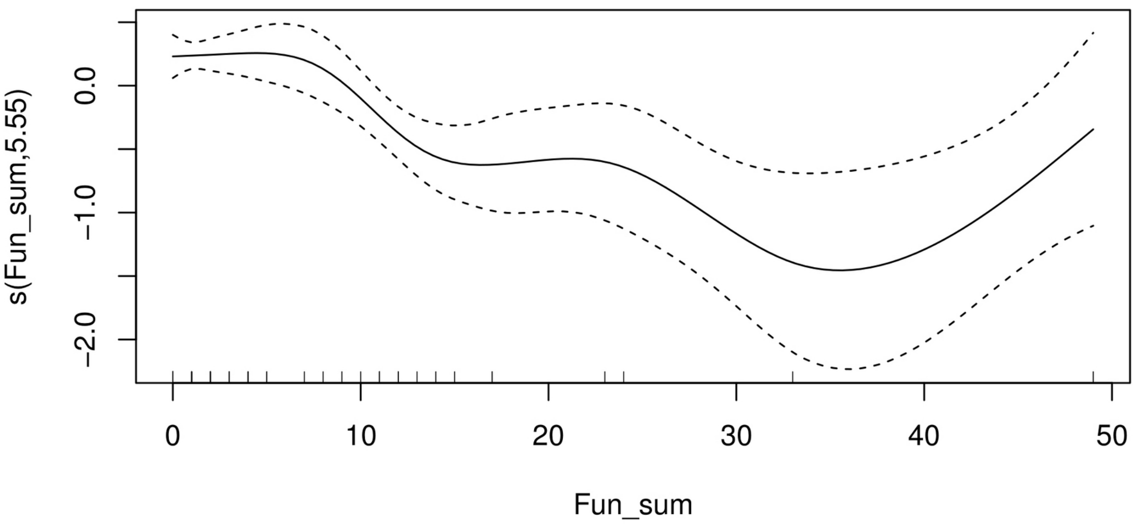

Figure S5. Abundance of raw reads affiliated to 7 algal mOTUs which appear to be signature "species" for second perturbation period from 7.3 to 5.5 kyr (grey area). X68484- *Scherffelia dubia*, AY197623 – *Choricystis* sp. MDL1/12-8, FN298929 - *Choricystis* sp. TP-2009, AB771902/LC425383 - *Desmodesmus costato-granulatus*, AY919716/EF058253- *Woloszynskia pascheri*, AY195976 - *Gloeotila* sp. JL11-10, DQ388552 – *Spumella vulgaris*

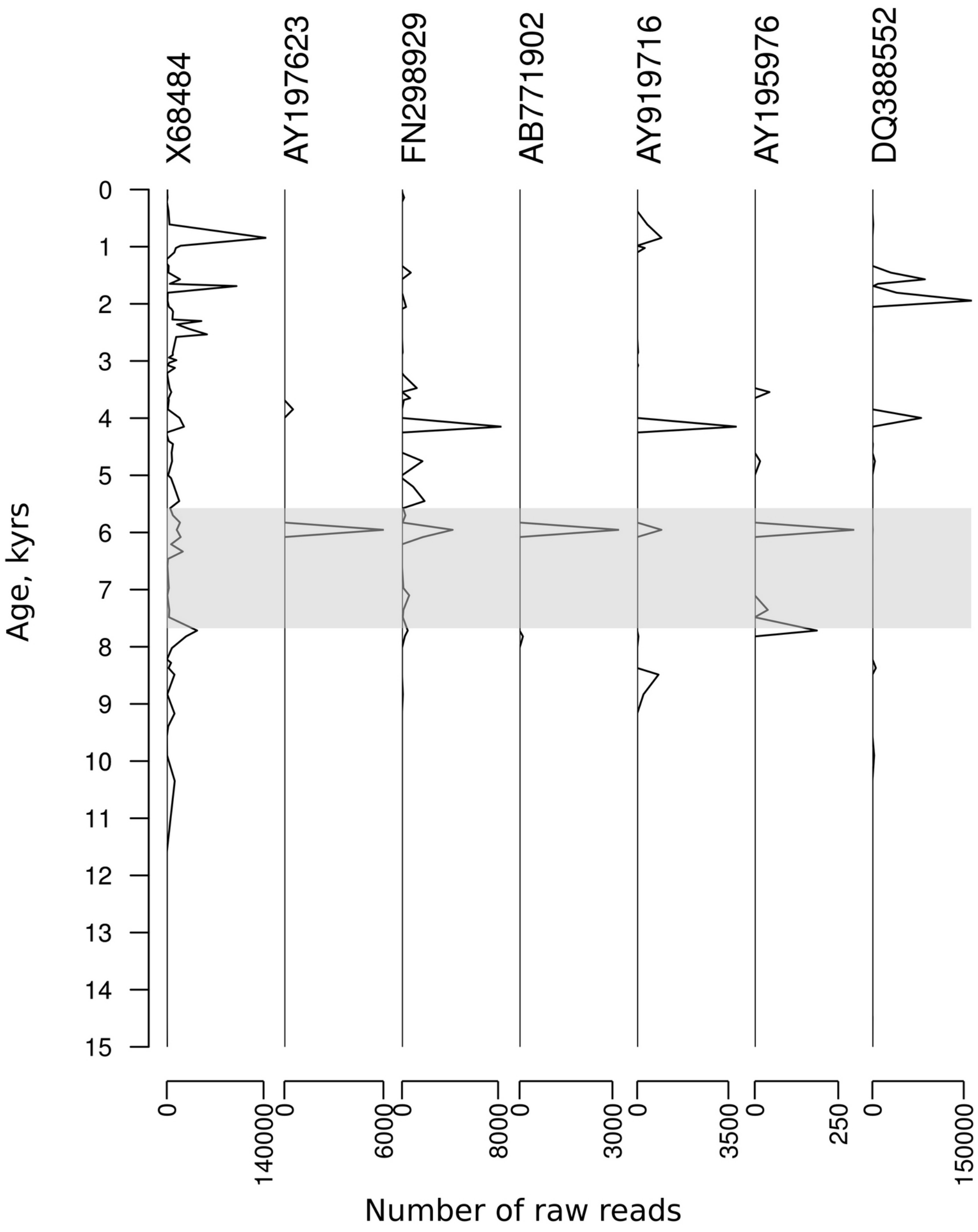
